# Effects of temperature on Zika dynamics and control

**DOI:** 10.1101/855072

**Authors:** Calistus N Ngonghala, Sadie J. Ryan, Blanka Tesla, Leah R. Demakovskys, Erin A Mordecai, Courtney C. Murdock, Matthew H. Bonds

## Abstract

When a formerly rare pathogen emerges to cause a pandemic, it is critical to understand the ecology of the disease dynamics and its potential effects on disease control. Here, we take advantage of newly available experimental data to parameterize a temperature-dependent dynamical model of Zika virus (ZIKV) transmission, and analyze the effects of temperature variability and the parameters related to control strategies on ZIKV *R*_0_ and the final epidemic size (i.e., total number of human cases). Sensitivity analyses identified that *R*_0_ and the final epidemic size were largely driven by different parameters, with the exception of temperature, which is the dominant driver of epidemic dynamics in the models. Our estimate of *R*_0_ had a single optimum temperature (≈ 30° C), comparable to recently published results (≈ 29°)^1^. However, the total number of human cases (“final epidemic size”) is maximized across a wider temperature range, from 24 to 36°C. The models indicate that the disease is highly sensitive to seasonal temperature variation. For example, although the model predicts that Zika transmission cannot occur at a constant temperature of 22°C, with seasonal variation of 5°C around a mean of 22°C, the model predicts a larger epidemic than what would occur at a constant 30°C, the temperature predicted to maximize *R*_0_. This suggests that the potential geographic range of Zika is wider than indicated from static *R*_0_ models, underscoring the importance of climate dynamics and variation on emerging infectious diseases.

## Introduction

Vector-borne viruses (arboviruses) are emerging threats to both human and animal health. The global expansion of dengue virus (DENV), West Nile virus (WNV), chikungunya (CHIKV) and most recently Zika virus (ZIKV) are prominent examples of how quickly mosquito-transmitted viruses can emerge and spread through naive host populations. Currently 3.9 billion people living within 120 countries are at risk of mosquito-borne arboviral diseases^2^ with effects on human well-being that can be devastating (e.g., death, illness, as well as social and human ramifications of Zika induced-microcephaly and other congenital disorders)^3^. Anticipating and preventing outbreaks of emerging mosquito-borne viruses across these host populations is a major challenge.

Despite growing research to develop new therapeutics and vaccines, mitigating arbovirus disease spread still depends on conventional mosquito control methods, often with mixed success. Developing tools that allow us to successfully predict outbreaks of these viruses and efficiently target current and future interventions to specific times and locations can aid effective mosquito and disease control. Such efforts are often limited by gaps in knowledge on the relationships among mosquito vectors, pathogens, and the environment, especially for emerging arboviruses such as CHIKV and ZIKV. Even in well-researched disease systems (e.g. malaria and DENV), key transmission parameters are only estimated from a few studies^4–6^.

Variation in environmental temperature has a strong impact on the environmental suitability for transmission risk across a diversity of vector-borne disease systems^7–10^. Mosquitoes are small ectothermic organisms, and their fitness^11, 12^, life history^13–18^, and vectorial capacity^4–6, 17, 19–22^ exhibit non-linear, unimodal relationships with environmental temperature. Recent work by Tesla et al.^20^ demonstrates such temperature-transmission relationships for ZIKV, a recently emerging pathogen. These temperature-transmission relationships have significant ramifications on how disease transmission varies seasonally, across geographic locations, and with future climate and land use change. Control tools being considered for use within integrated vector management (IVM) strategies may also be affected by temperature, such as conventional chemical insecticides that target a diverse range of insect pests^23–28^, including mosquitoes^29, 30^. Further, there is evidence that temperature could modify the efficacy of novel control interventions, such as mosquito lines transinfected with the intracellular bacteria *Wolbachia*^31–34^.

Several modeling frameworks have been used to predict environmental suitability for vector-borne disease transmission, including, most recently, temperature-dependent *R*_0_ models^4–6, 19, 20^ and compartmental models of vector-borne disease dynamics^9, 35, 36^. The parameter *R*_0_ is broadly considered to be the most important summary statistic in epidemiology and disease ecology. It is defined as the expected number of new human (respectively, mosquito) cases generated by a single infectious human (respectively, mosquito) introduced into a fully susceptible human (respectively, mosquito) population throughout the period within which that human (respectively, mosquito) is infectious^37^. As a simple metric, it can easily incorporate the non-linear influence of multiple temperature-dependent mosquito and pathogen traits, and has been applied to define the thermal optimum and limits for malaria^5, 6, 38^, DENV, CHIKV^4, 39, 40^, ZIKV^4, 20^, and Ross River virus^19^. However, temperature-dependent *R*_0_ formulations only define the relative risk of disease emergence and do not predict the final epidemic size (or incidence). The derivation, interpretation, and validation of *R*_0_ models is thus problematic in highly variable systems^41^. Dynamical models of transmission that track densities of infectious individuals over time, on the other hand, can more readily capture the impact of varying environmental conditions.

To better understand potential climate effects on control strategies for ZIKV, we developed a temperature-dependent dynamical model based on recent experimental work characterizing temperature-trait relationships between ZIKV vector competence, extrinsic incubation rate, and the per capita daily mosquito mortality rate^20^. Through numerical and sensitivity analyses we analyze effects of control parameters and temperature on *R*_0_ and the final epidemic size. The model addresses the following questions: 1) How do the thermal optima and ranges for *R*_0_ compare to those for the human final epidemic size? 2) How does seasonal temperature variation affect the final epidemic size relative to a constant temperature environment? 3) Which parameters have the greatest impact on *R*_0_ and the final epidemic size that can inform control efforts? 4) Are different thermal environments more or less suitable for specific control strategies?

Our results show that *R*_0_ and the final epidemic size were largely driven by different parameters, with the exception of temperature being the dominant driver of both transmission metrics. Further, the human final epidemic size was maximized across a wider range of temperatures than what would have been predicted from the temperature-dependent *R*_0_ model. The human final epidemic size was highly sensitive to seasonal temperature variation, suggesting the potential invasion map of ZIKV may be wider than previously reported. Further, the effectiveness of potential control strategies (e.g., vaccines, drug treatment, and insecticides) are predicted to be sensitive to such differences in seasonal temperature variation.

## Methods

We construct a temperature-dependent compartmentalized model of ZIKV dynamics with and without seasonal temperature change. Where possible, model parameters are estimated from the most recent laboratory experiments on temperature effects on the life cycle of the virus^4, 42^. We compare how temperature dependence affects *R*_0_, the human “final epidemic size”(total number of infected individuals over the course of the epidemic), and key ecological characteristics of the system, such as extrinsic incubation period, the probability of transmission from the mosquito to the human, the probability of transmission from the human to the mosquito, and daily rates of mosquito and egg to adult survival. We then analyze the combined effects of disease control parameters and temperature on *R*_0_ and the final epidemic size. Through a Latin-Hypercube Sampling-based^43^ sensitivity analysis we identify key parameters that most drive the epidemiological outcomes (*R*_0_ and the final epidemic size).

### The basic dynamic model

The model apportions humans into four groups based on ZIKV infection status that changes over time, *t*: susceptible *S*_*h*_ (not infected), latent *E*_*h*_ (contracted the virus, but not yet infectious), infectious *I*_*h*_ (contracted the virus and can transmit it), and recovered *R*_*h*_ with lifelong immunity. The mosquito population is divided into similar classes, where the state variables have subscript *v*, but without an immune class since it is assumed that infectious mosquitoes do not clear the virus once it is in the salivary glands. The total human and mosquito populations are *N*_*h*_ = *S*_*h*_ + *E*_*h*_ + *I*_*h*_ + *R*_*h*_ and *N*_*v*_ = *S*_*v*_ + *E*_*v*_ + *I*_*v*_.

We assume that the human population is constant during an outbreak (relevant for short epidemics). Susceptible humans acquire the virus at rate (force of infection) 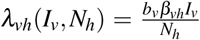, while susceptible mosquitoes acquire the virus at rate 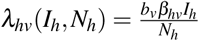, where *b*_*v*_ is the number of human bites per mosquito per unit time, *β*_*vh*_ is the probability that an infectious mosquito successfully transmits the virus while taking a blood meal from a susceptible human (i.e., the transmission rate), and *β*_*hv*_ is the probability that an infectious human successfully transmits the virus to a biting, susceptible mosquito (i.e. the infection rate). The respective average residence times of infected humans and mosquitoes in the latent classes are 1*/σ*_*h*_ and 1*/σ*_*v*_, while the respective rates at which humans and mosquitoes become infectious are *σ*_*h*_ and *σ*_*v*_. Humans are infectious for approximately 1*/γ*_*h*_ days before recovering with permanent immunity (*γ*_*h*_ is the per capita human recovery rate), while infectious mosquitoes remain infectious until they die. Mosquito recruitment occurs at a per capita rate 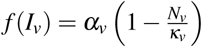, where *κ*_*v*_ is the carrying capacity (maximum number of mosquitoes a breeding site can support). Further, *α*_*v*_ = *θ*_*v*_*ν*_*v*_*ϕ*_*v*_*/µ*_*v*_, consistes of *θ*_*v*_, or the number of eggs a female mosquito produces per day; *ν*_*v*_, the probability of surviving from egg to adult; and *ϕ*_*v*_, the rate at which an egg develops into an adult mosquito. Mosquitoes die naturally at per capita rate *µ*_*v*_, where 1*/µ*_*v*_ is the average lifespan of mosquitoes. See Fig. 1 for a schematic of the model and Table 1 for details on parameter values. The dynamic model for the Zika virus is described by the equations:

**Table 1.**
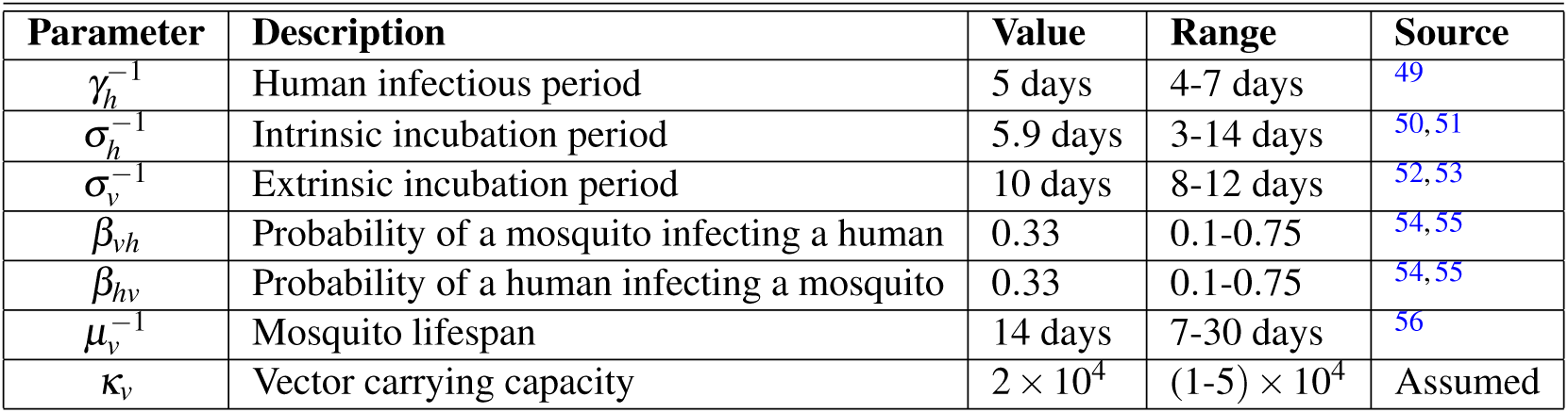
Parameter (P) definitions and baseline values for system without temperature dependence.

**Figure 1.**
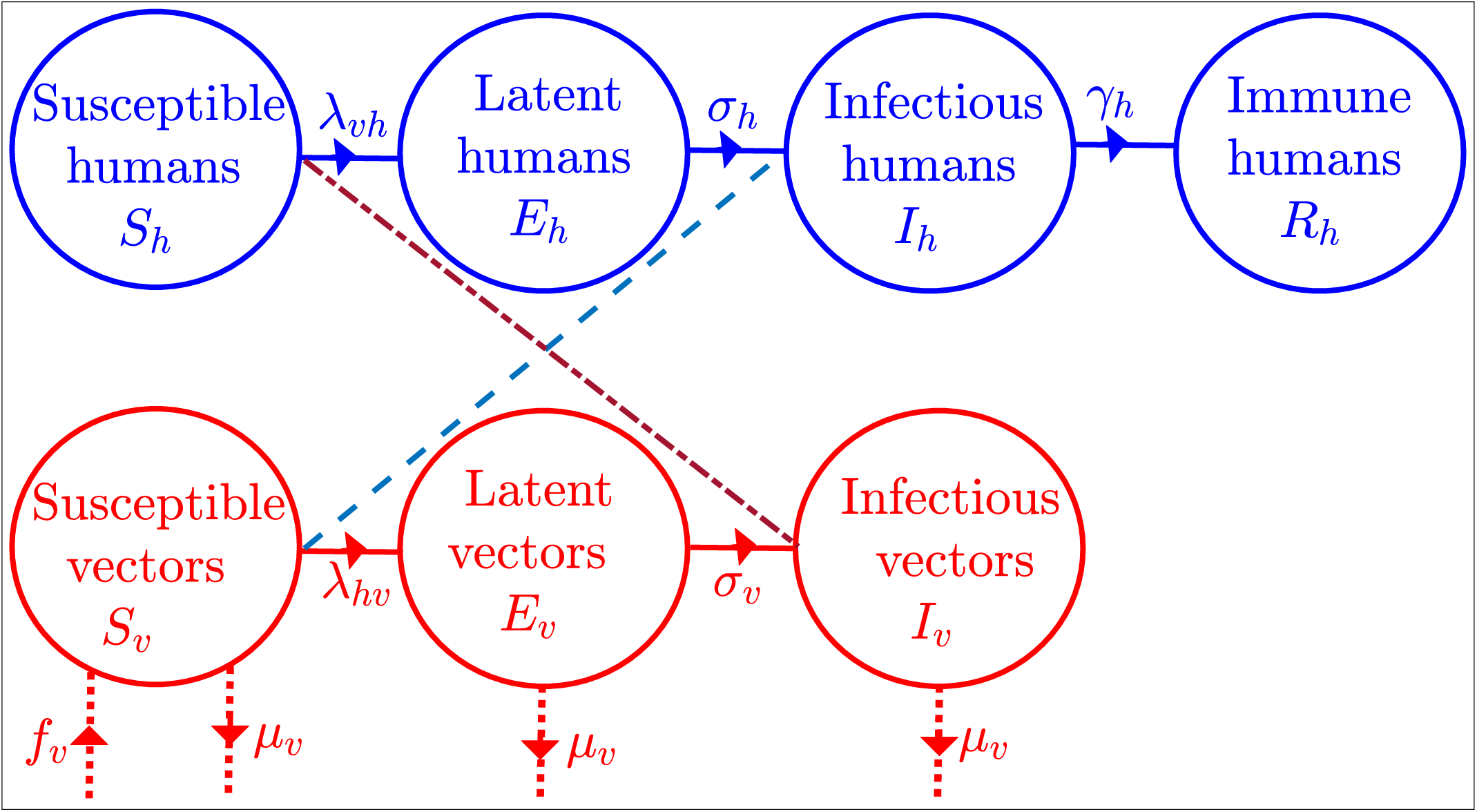
Compartmental model of Zika virus transmission. Compartments are divided into humans (blue), and vectors (red), representing disease status, with transitions between compartments (rates) in solid lines. The transmission of Zika virus from humans to vectors is denoted by dashed lines, and from vectors to humans by dashed-dotted line. Rates of demographic change (births and deaths) in the vector population are denoted by dotted lines.

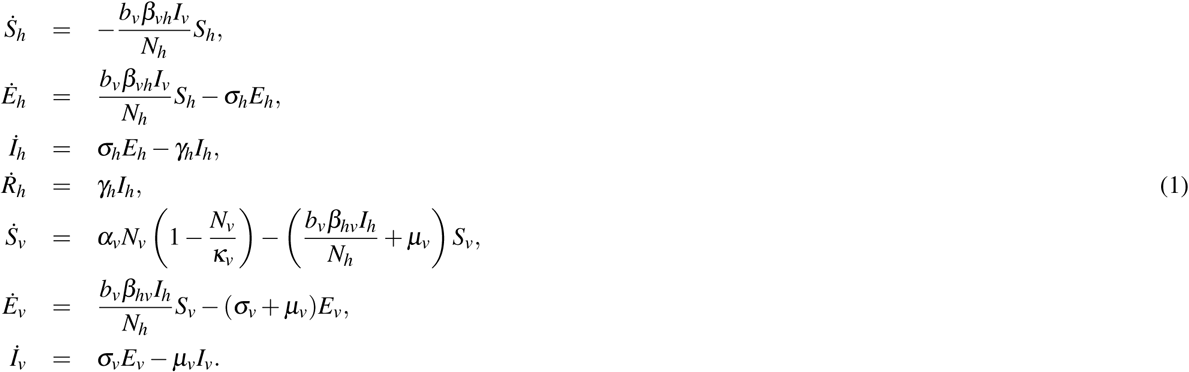

Dots denote differentiation with respect to time, *t*. The dynamics of the total human population and mosquito populations are described by the equations:

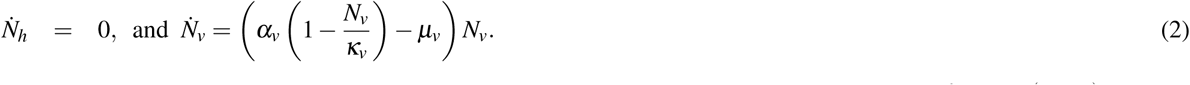

Without Zika virus, the mosquito population grows according to Eq. (2), or 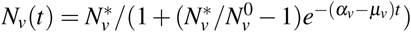, where 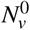 is the initial mosquito population and 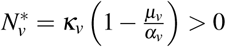 for *α*_*v*_ > *µ*_*v*_ is the positive equilibrium obtained by setting the right-hand-side of the equation to zero. Observe that 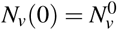, and that when *α*_*v*_ > *µ*_*v*_, the total mosquito population relaxes on the equilibrium population 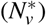 in the long-run. Therefore, the equilibrium point 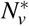 is stable when *α*_*v*_ > *µ*_*v*_ and vanishes when *α*_*v*_ < *µ*_*v*_. The case for which *α*_*v*_ < *µ*_*v*_ results in a trivial mosquito equilibrium represents a situation in which the mosquito population becomes extinct.

In the presence of the Zika virus, the basic reproduction number of system (1) is:

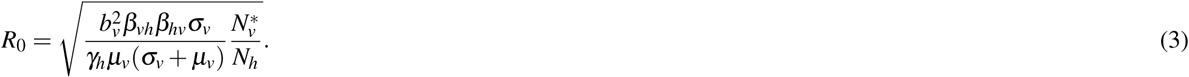

The main difference between this *R*_0_ calculation and that from the Ross-MacDonald model is in the probability that the mosquito survives the latent period. The Zika virus can spread when *R*_0_ > 1 and can be contained when *R*_0_ < 1.

For the purposes of exploring control strategies, we also consider a variant of this model that includes vaccination, where vaccinated susceptible humans are assumed to enter the immune class directly. For this special case, the first and fourth equations of (1) are replaced by 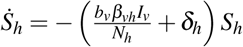 and 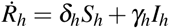, respectively, where *δ*_*h*_ represents the per capita rate human vaccination rate.

### Introducing temperature

The majority of the parameters associated with the mosquito vector (*θ*_*ev*_, *ν*_*v*_, *ϕ*_*v*_, *b*_*v*_, *µ*_*v*_), as well as ZIKV transmission (*β*_*hv*_) and replication (*σ*_*v*_), are known to be influenced by environmental and climate conditions^1, 4^. We investigate the effects of temperature variation on the dynamics of the mosquito population and ZIKV transmission over time. We follow the approach in^9^ and model temperature-dependent parameters with the functional forms presented in Table 2. We rely on values and ranges of temperature-dependent parameters from recent laboratory-generated analyses for Zika virus^1^ and *Ae. aegypti* life history parameters (e.g., the biting rate of mosquitoes, the number of eggs a female mosquito lays per day, the probability of an egg surviving to an adult mosquito, and the rate at which an egg develops into an adult mosquito) from^4^ as specified in Table 2. As in^4^, the functional forms for the temperature dependent parameters are based on the quadratic and Briere^44^ forms.

**Table 2.** Parameter values for temperature-dependent functional forms. The temperature in degrees Celsius, the minimum temperature, and the maximum temperature are denoted by *T, T*_0_, and *T*_*m*_, respectively. The parameter *c*, is a rate scaling factor, and the subscripts denote the corresponding traits, e.g., 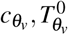 and 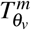 are for the number of eggs laid by a female mosquito per day, *θ*_*v*_. See Eqs. (4) for brief descriptions of the temperature-dependent parameters.

| | $c$ | | $T^0$ | | $T^m$ | | |
| --- | --- | --- | --- | --- | --- | --- | --- |
| Trait | Mean | Range | Mean | Range | Mean | Range | Source |
| $\theta_v(T)$ | $8.56 \times 10^{-3}$ | $[3.78, 14.1] \times 10^{-3}$ | 14.58 | [8, 08, 20.60] | 34.61 | [34, 35.77] | 4 |
| $v_v(T)$ | $-5.99 \times 10^{-3}$ | $[-6.82, -5.13] \times 10^{-3}$ | 13.56 | [12.56, 14.51] | 38.29 | [38.29, 39.02] | 4 |
| $\phi_v(T)$ | $7.86 \times 10^{-5}$ | $[5.75, 9.93] \times 10^{-5}$ | 11.36 | [7.19, 15.03] | 39.17 | [39.17, 39.54] | 4 |
| $b_v(T)$ | $2.02 \times 10^{-4}$ | $[1.2, 2.8] \times 10^{-4}$ | 13.35 | [5.84, 14.82] | 40.08 | [36.60, 40.51] | 4 |
| $\beta_{hv}(T)$ | $-3.54 \times 10^{-3}$ | $[-5.6, -1.8] \times 10^{-3}$ | 22.72 | [21.09, 24] | 38.38 | [36.46, 40.25] | 1 |
| $\sigma_v(T)$ | $1.74 \times 10^{-4}$ | $[5.4, 30.4] \times 10^{-5}$ | 18.27 | [7.68, 24] | 42.31 | [39.26, 45] | 1 |
| $1/\mu_v(T)$ | $-3.02 \times 10^{-1}$ | $[-4.68, -1.34] \times 10^{-1}$ | 11.25 | [6.3, 15.06] | 37.22 | [34.79, 39.57] | 1 |

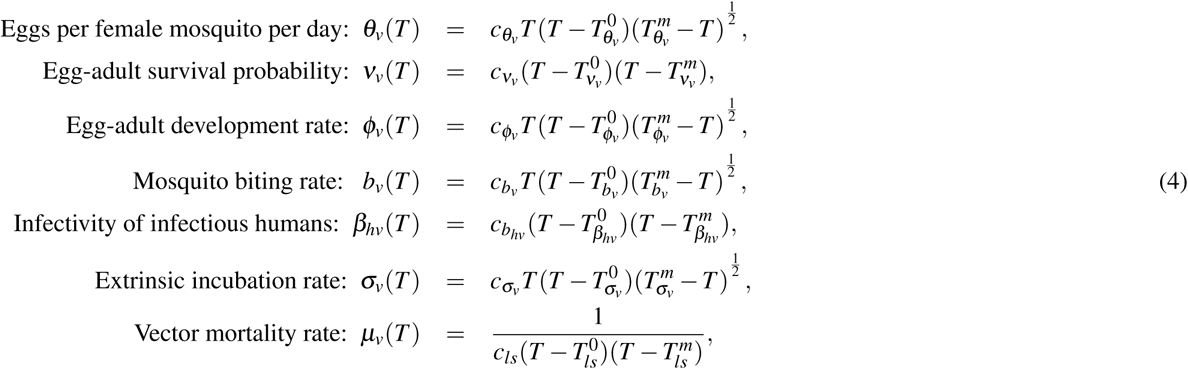

These relationships between some model parameters, model outcomes, and temperature are illustrated in Fig. 2. We further introduce seasonal variation in the system by modeling temperature through the functional form:

**Figure 2.**
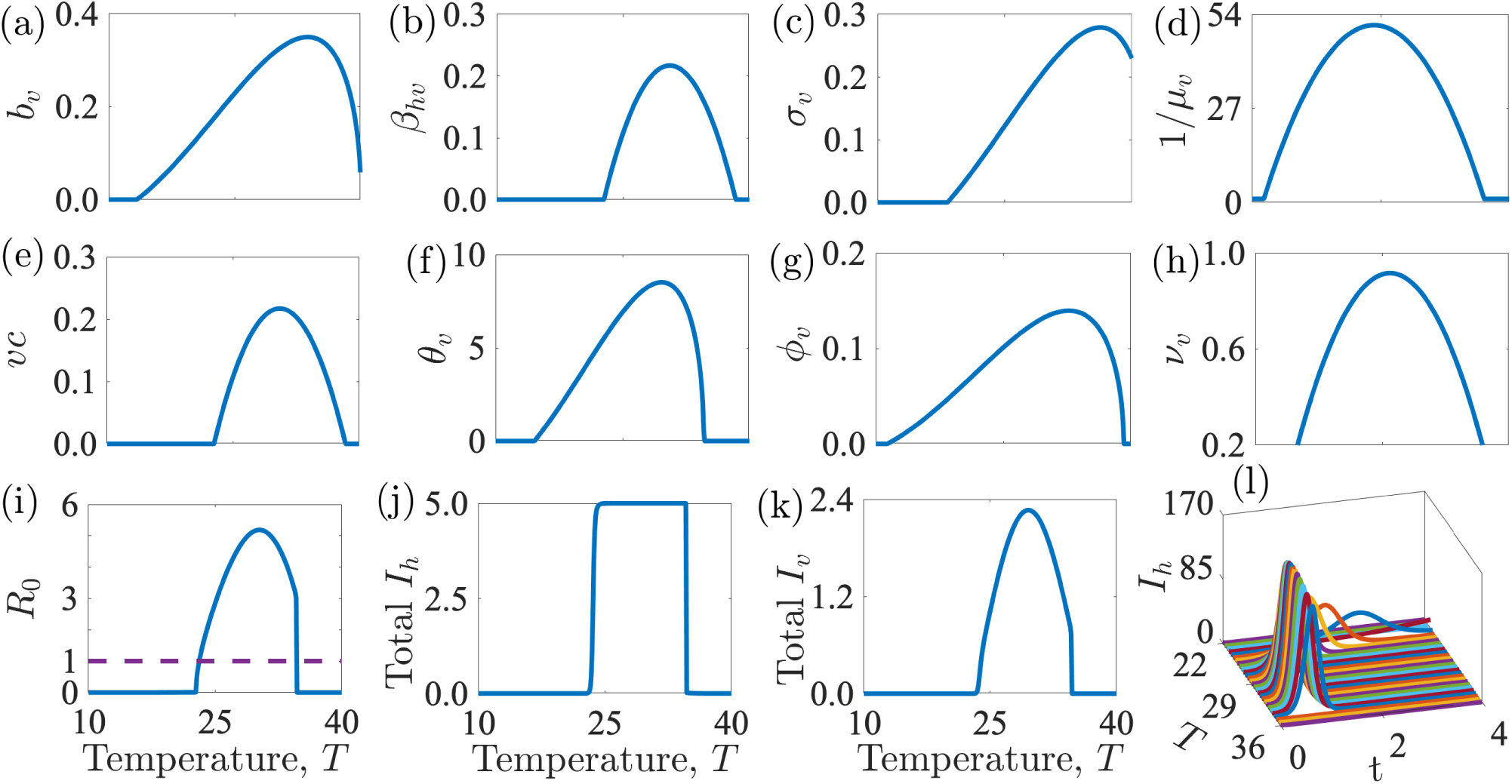
Effect of temperature, *T* on mosquito and pathogen parameters: (a) mosquito biting rate, *b*_*v*_; (b) the pathogen transmission probability from humans to mosquitoes, *β*_*hv*_; (c) the inverse of the pathogen extrinsic incubation period, *σ*_*v*_, (d) the average mosquito lifespan, 1*/µ*_*v*_; (e) the vector competence, *vc* = *β*_*hv*_*β*_*vh*_; (f) the number of eggs laid by a female mosquito per day, *θ*_*v*_; (g) the egg-to-adult mosquito development rate, *ϕ*_*v*_; (h) the egg-to-adult mosquito survival probability, *ν*_*v*_; (i) the basic reproduction number, *R*_0_; (j and k) the total infectious human population, *I*_*h*_ in thousands, and total vector population, *I*_*v*_ in hundreds of thousands; and (l) the infectious human population. Time, *t* in (l) is in hundreds of days. The number of infectious individuals rises with temperature up to an optimal temperature between 29°C and 32°C. As temperatures are increased beyond the optimal, the number of infectious individuals falls.

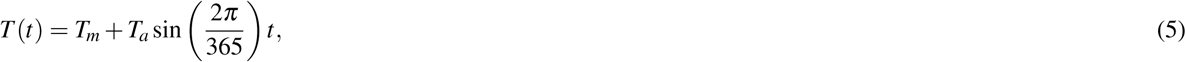

where *T*_*m*_ is the mean annual temperature and *T*_*a*_ is the amplitude (divergence from mean temperature or mid-point between the lowest and highest annual temperatures). Temperature-dependent parameters used in our analyses are presented in Table 2.

### Control strategies

To analyze the relationship between temperature and Zika control, we identified the following parameters of the system that correspond to potential control strategies: vaccination (*δ*_*h*_) decreases susceptibility and is directly incorporated into the models as described above; recovery rates (*γ*_*h*_) can, for example, be increased through treatment with antiviral medication; vector biting rates (*b*_*v*_) can be reduced through decreasing exposure to mosquitoes with personal protection or household improvements; vector-to-human transmission probability (*β*_*vh*_) can decrease with transmission-blocking *Wolbachia*; the vector carrying capacity (*κ*_*v*_) can be reduced by eliminating vector breeding grounds near human habitats; egg-adult survival probability (*ν*_*v*_) can be reduced through larvicides; and adult mosquito survival rate (*µ*_*v*_) can be decreased through indoor spraying and the use of aldulticides. We investigate how the interactions of these control parameters and temperature influence *R*_0_ and the final epidemic size, total *I*_*h*_.

### Sensitivity analysis

Two types of sensitivity analyses - local and global - were used to explore the impact of temperature and selected control parameters on the basic reproduction number (*R*_0_) and the human final epidemic size (total *I*_*h*_). The local sensitivity analysis was conducted by varying only one parameter while holding all other parameters fixed, or varying both temperature and a control parameter while holding the other parameters fixed. Each varied parameter was divided into 50, 100, and 250 equally spaced points within biologically feasible bounds. See Figs. 2-6 for results. As the human vaccination rate (*δ*_*h*_) does not appear explicitly in the expression of *R*_0_, we cannot assess the impact of temperature on this control parameter on the basic reproduction number. However, we explore the impact of temperature on the human vaccination rate and associated implications for *I*_*h*_, the human final epidemic size (Fig. 3).

**Figure 3.**
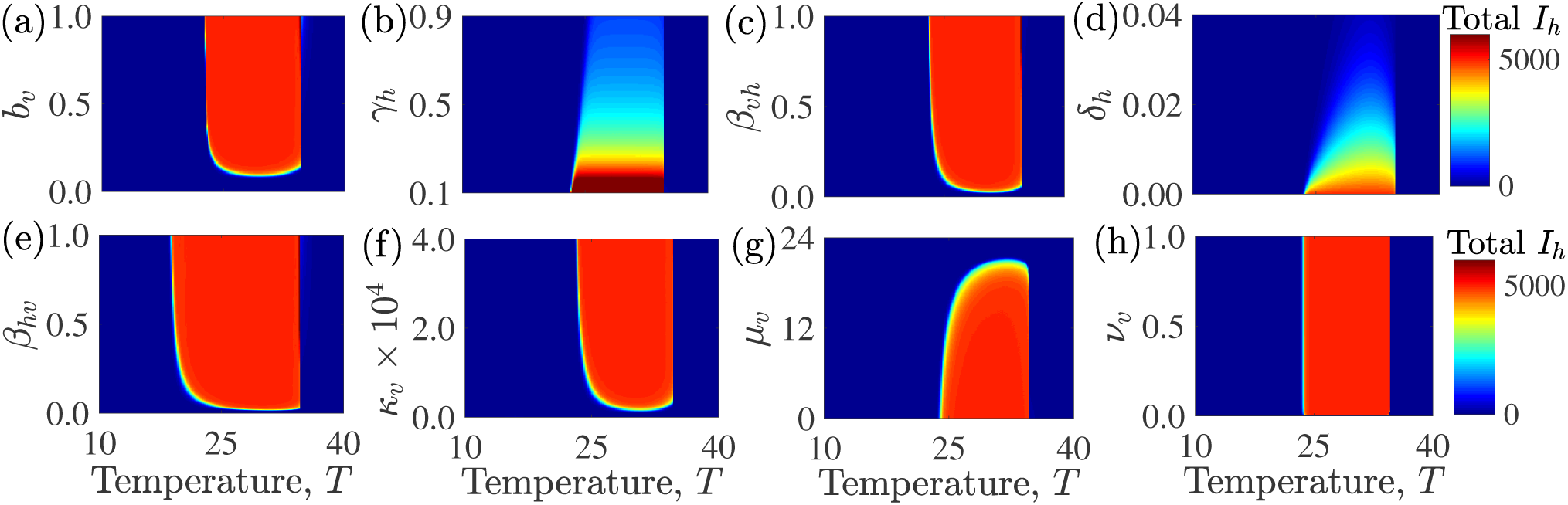
Effect of temperature and control parameters on human the final epidemic size (total infectious human population): (a) vector biting rate, *b*_*v*_; (b) human recovery rate, *γ*_*h*_; (c) human vaccination rate, *δ*_*h*_; (d) the probability of transmission from the mosquito to the human, *β*_*vh*_; (e) the probability of transmission from the human to the mosquito, *β*_*hv*_; (f) vector mortality rate, *µ*_*v*_; (g) vector carrying capacity, *κ*_*v*_; (h) and the vector egg-adult survival probability, *ν*_*v*_. With the exception of the human vaccination and recovery rates, temperature does not substantially alter the effects of most control parameters on the human final epidemic size (the color bands have little gradient in the parameter space).

**Figure 4.**
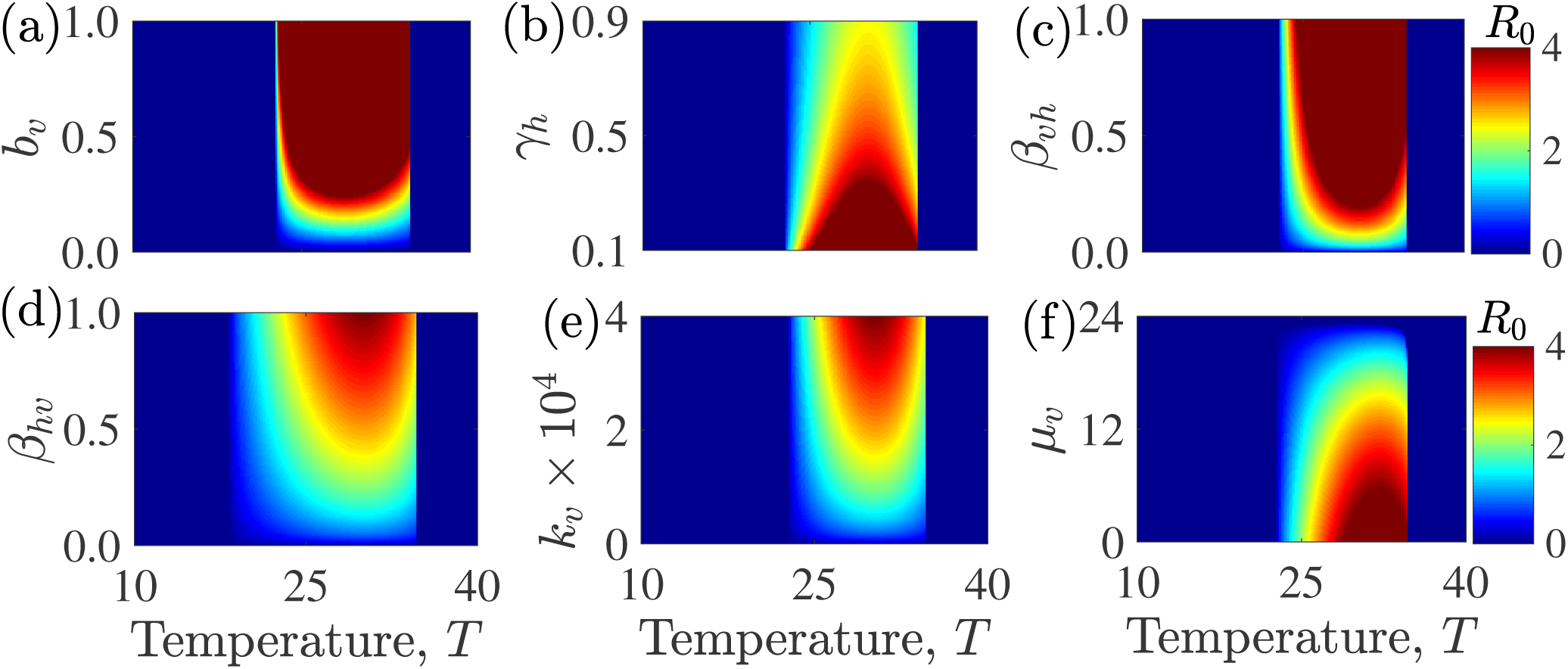
Effect on the basic reproduction number, *R*_0_, of temperature and control parameters: (a) the mosquito biting rate, *b*_*v*_; (b) the human recovery rate, *γ*_*h*_; (c) the probability of transmission from the mosquito to the human, *β*_*vh*_; (d) the probability of transmission from the human to the mosquito, *β*_*hv*_; (e) the mosquito carrying capacity, *κ*_*v*_; and (f) the mosquito mortality rate, *µ*_*v*_. (Vaccination *δ*_*h*_ and the egg-to-adult mosquito survival probability, *ν*_*v*_, do not appear explicitly the in *R*_0_ model.) The effect of these parameters on *R*_0_ is more dependent on temperature than their effects on the final epidemic size (e.g., the color bands are less vertical and more diagonal in parts of the parameter space).

**Figure 5.**
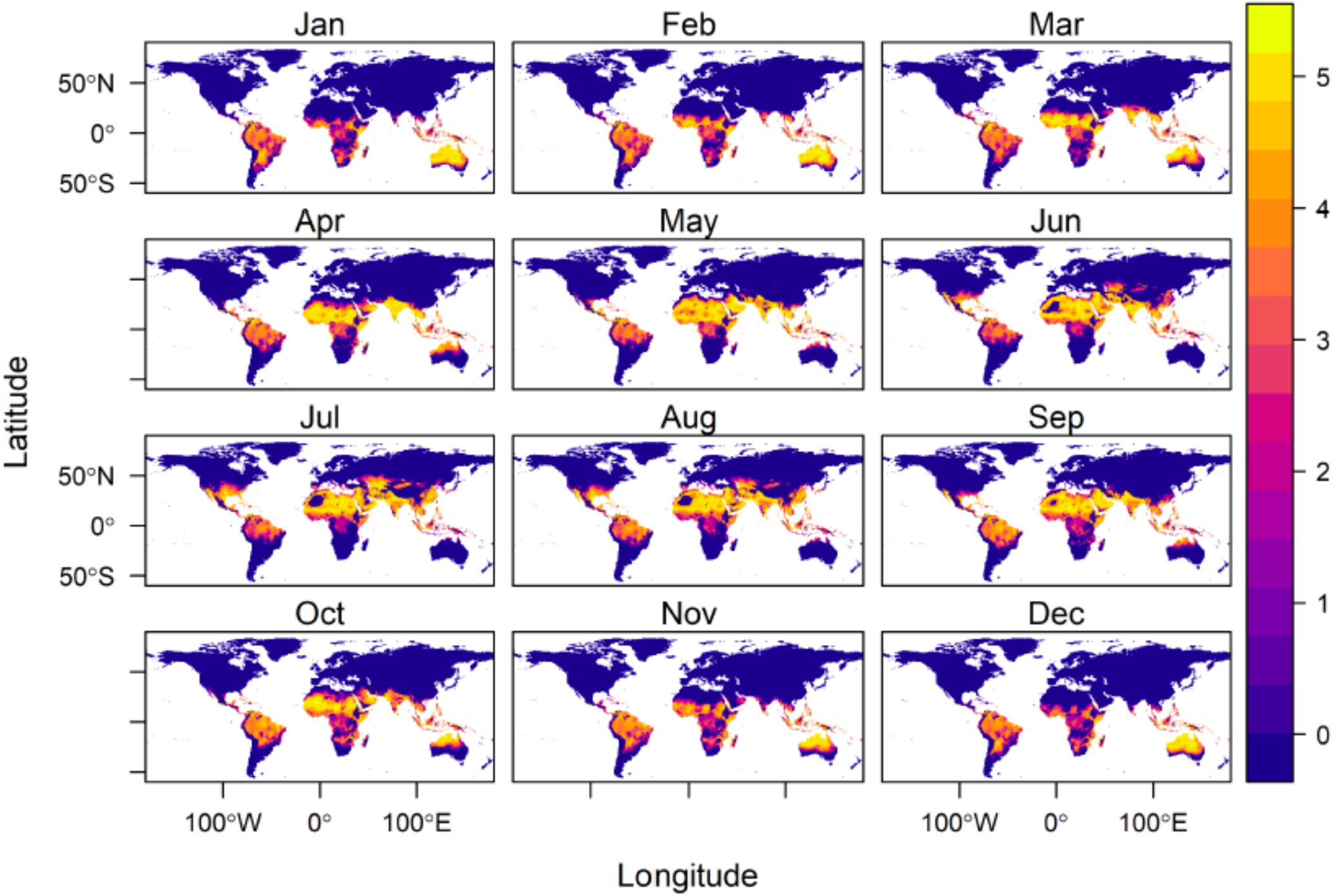
Estimated basic reproduction number *R*_0_, for different mean monthly temperatures, globally. The color bar represents increasing *R*_0_ values with the lowest values denoted by dark blue and the highest values denoted by light yellow. A value of *R*_0_ below 1 means the disease cannot take off. Note the month-to-month variation in locations such as Australia, demonstrating the impact of seasonal temperature variation on *R*_0_, and therefore on the control level needed to reduce or eradicate disease.

**Figure 6.**
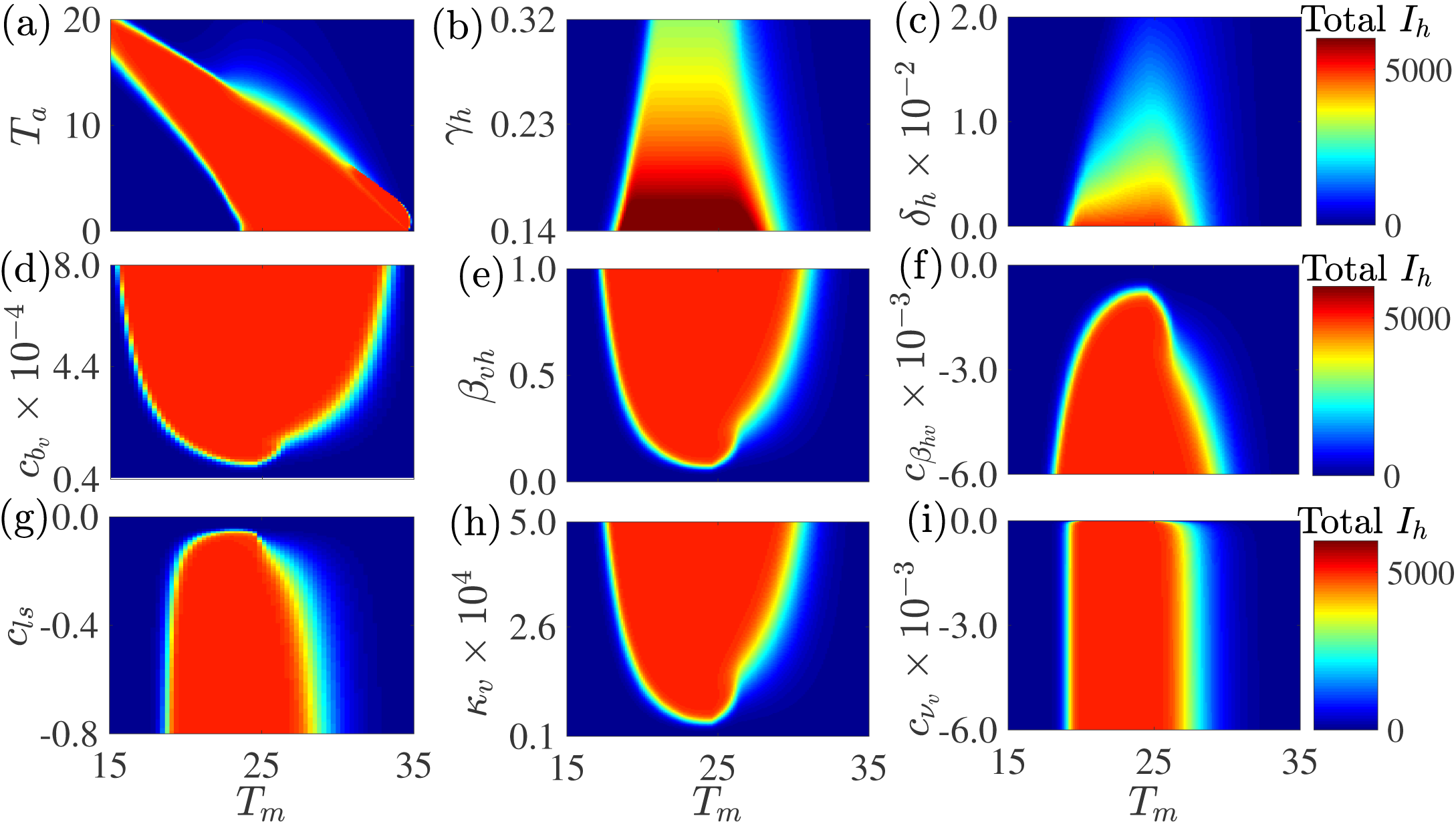
The human final epidemic size is sensitive to the annual mean (oscillation) temperature, *T*_*m*_, and several important control-related parameters: (a) the seasonal divergence of the annual temperature from the mean, *T*_*a*_; (b) the human recovery rate, *γ*_*h*_; (c) the human vaccination rate, *δ*_*h*_; (d) the scaling factor of vector biting rate, *c*_*b_v_*_; (e) the probability of transmission from the mosquito to the human, *β*_*vh*_; (f) the scaling factor of the probability of transmission from the human to the mosauito, *c*_*β_hv_*_; (g) the scaling factor of the vector mortality rate, *c*_*ls*_ (g); (h) the vector carrying capacity, *κ*_*v*_; (i) and the egg survival probability scaling factor, *c*_*ν_v_*_. The annual mean (oscillation) temperature varies along the x-axis, the seasonal divergence of the annual temperature from the mean and the other control parameters vary along the y-axes, and the color scale indicates the total infectious humans. Apart from (a), where the temperature amplitude (*T*_*a*_) is varying, the amplitude is set at 10° C for the other plots. Plot (a) shows that there can be large epidemics even when mean temperatures are low if the seasonal variation (the amplitude) is high enough, as would be found in subtropical and temperate regions.

Global sensitivity analysis is presented in Fig. 7. The analysis is carried out using the Latin-hypercube Sampling (LHS) and Partial Rank Correlation Coefficient (PRCC) technique^43^. The process involves identifying a biologically feasible mean, minimum and maximum value for each of the parameters (see, for example,^1, 4^) and subdividing the range of each parameter into 1000 equal sub-intervals, assuming a uniform distribution between the minimum and maximum values of each parameter. We then sample at random and without replacement from the parameter distributions to generate an *m* × *n* latin-hypercube sampling matrix, whose *m* rows (i.e., 1000 rows) consist of different values for each of the model parameters and the *n* columns (corresponding to the number of parameters in the system) consist of different values for the same parameter. Thus, each row of the Latin hypercube sampling matrix provides a parameter regime that is used for computing the basic reproduction number, solving the dynamic system, and computing the human final epidemic size. The parameters, basic reproduction number, and the human final epidemic size are then ranked with partial correlation coefficients estimated for each parameter along with corresponding p-values. PRCCs range from − 1 to 1 and are used to examine the correlation between model parameters and model outputs (*R*_0_ and the final epidemic size). This method thus identifies parameters with the most significant influence on model outputs; it does not quantify the effect of a change in a parameter on the output.

**Figure 7.**
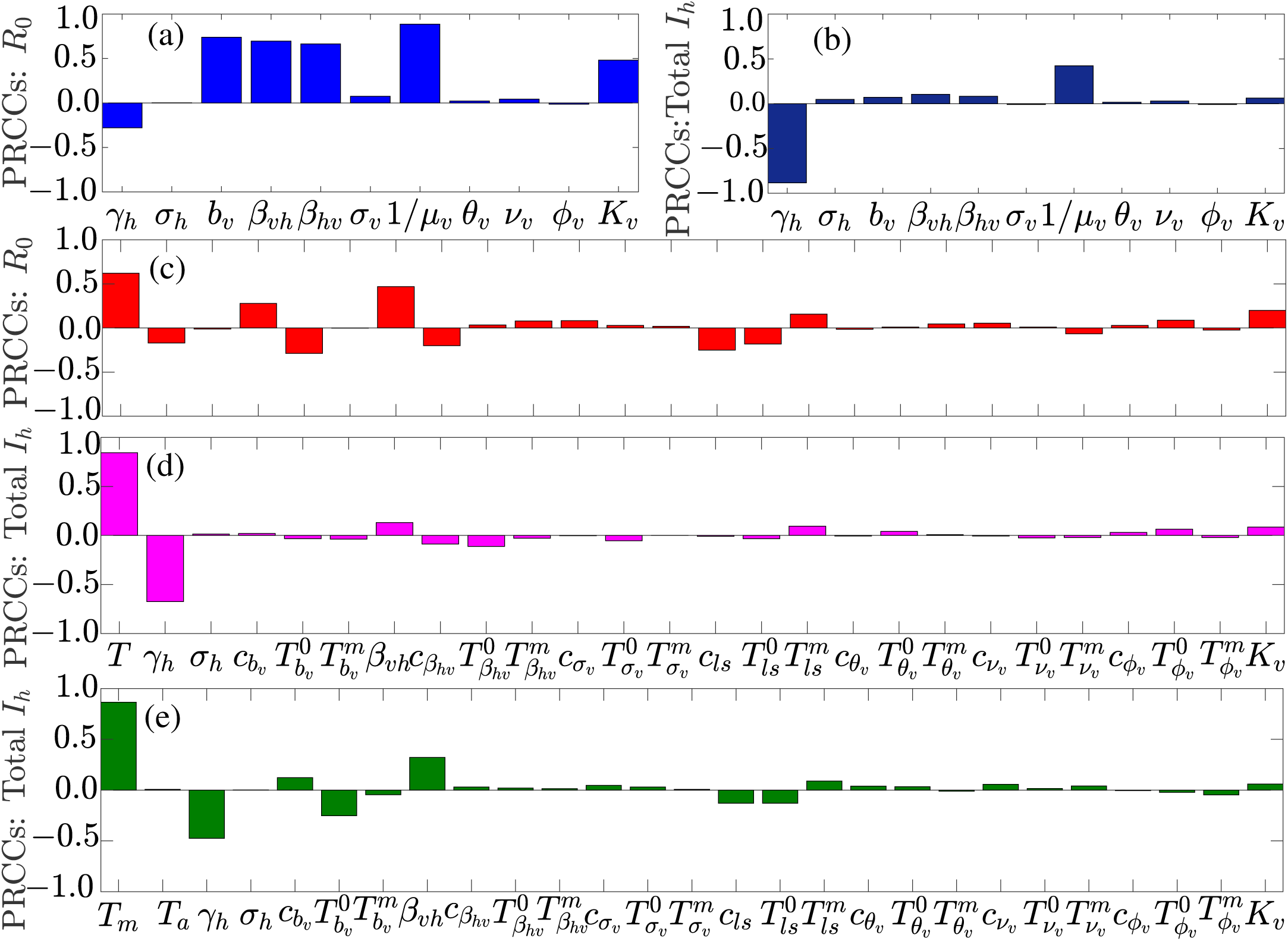
Global sensitivity analysis indicates the sensitivity of the basic reproduction number, *R*_0_ (a and c), and the final epidemic size, total *I*_*h*_ (b, d, and e), to all model parameters. Bars indicate partial rank correlation coefficients (PRCC), illustrating the contribution of parameters to variability or uncertainty in the model outputs (*R*_0_ and *I*_*h*_). (a)-(b) No temperature dependence; (b)-(c) temperature dependence but no temperature variation; (e) temperature dependence and temperature variation. Without accounting for temperature, models show different sets of drivers for *R*_0_ (a) than burden (b). When temperature is included, it is the dominant contributor to model sensitivity for all models (c)-(e).

### Mapping Seasonal Control

We mapped the *R*_0_ as a function of monthly mean temperature (Fig. 5). Globally gridded monthly mean current temperature were downloaded from WorldClim.org^45^, at a 5 arc-minute resolution (approximately 10 *km*^2^ at the equator), and predicted rates as a function of temperature at 0.2^0^ *C* were mapped to the global grids. All raster calculations and graphics were conducted in R, using package raster^46^.

## Results

### Impact of temperature on model parameters and key outputs

The models show unimodal relationships between temperature and the temperature-dependent parameters, resulting in an optimal temperature that maximizes parameter values and a critical minimum and maximum temperature at which parameter values go to zero. Figure 2 presents the effects of temperature on mosquito and pathogen parameters, the final epidemic size in humans (total number of infected individuals over the course of the epidemic) and mosquitoes, and the basic reproduction number, *R*_0_ (via temperature effects on mosquito and ZIKV parameters).

The response of *R*_0_ to temperature is strongly peaked as has been demonstrated in other systems (e.g., dengue, malaria, Ross River virus)^4, 42^. In contrast, the relationship between the final epidemic size and temperature discretely changes when temperature enters into a thermal range, but has little or no change to temperature variation within that range (Fig. 2 (i) versus (j)). At temperatures associated with lower epidemic peaks, there are longer epidemic periods, resulting in the same total number of infected individuals over the course of the epidemic (Figure 2 (l)).

### Zika virus control

For most control-related parameters, effects on the final epidemic size were largely independent of temperature (color bands are vertical for most of the plots, (Fig. 3(a-h)). However, vaccination and recovery were less influential on the final epidemic size at optimal temperatures, indicating a higher proportion of the population needs to be vaccinated at optimal temperatures than at sub-optimal temperatures to achieve a given reduction in the overall final epidemic size (Fig. 3(c)). Thus, warming temperatures (for most countries) will generally make it more difficult to control Zika through vaccination and drugs.

The effects on the basic reproduction number (*R*_0_) of most control parameters were more dependent on temperature (Fig. 4) than was the case for the final epidemic size (total *I*_*h*_), with the greatest effect involving the clearance rate of infection (*γ*_*h*_), the probability of transmission from an infectious human to a susceptible mosquito (*β*_*hv*_), mosquito carrying capacity (*κ*_*v*_), and the mosquito mortality rate (*µ*_*v*_).

### Seasonal variation

Seasonal temperature variation affects outcomes by providing transient temperatures (variation from the mean) where the basic reproduction number changes and can rise above 1 allowing for transmission to occur. At constant temperatures, epidemics only occur in humans between 23^0^ - 37^0^ C. Seasonal temperature variation of *±* 6^0^ C allows large epidemics to occur between mean temperatures of 15 - 34^0^ C (Fig. 6 (a)). Thus temperate regions with large seasonal variation can support large epidemics, comparable to that of warmer tropical climates with less seasonal variation. In contrast to the models without seasonal variation (Fig. 3), the models with seasonal variation (Fig. 6) indicate that the effectiveness of control parameters on the final epidemic size is generally sensitive to changes in temperature (e.g., the color bands in the subplots of Fig. 6 are diagonal in more of the parameter space than they are in (Fig. 3). Fig. 5 shows how the thermal conditions that are suitable for Zika (where *R*_0_ > 1) change with seasonal temperature variation across the globe.

### Global uncertainty and sensitivity analysis

A global sensitivity analysis using Latin Hypercube sampling showed that *R*_0_ and the final epidemic size are largely sensitive to different parameters. However, temperature is a dominant driver of variation in both the basic reproduction number (*R*_0_) and the final epidemic size (total *I*_*h*_) when it is included in the model (Fig. 7(c-e)). The human recovery rate, *γ*_*h*_, was a consistently influential driver of the final epidemic size. In contrast, the basic reproduction number was not sensitive to recovery in the models with and without temperature. While *R*_0_ was also sensitive to vector competence (*β*_*vh*_ and *β*_*hv*_), biting rate (*b*_*v*_), and mosquito lifespan (1/*µ*), total infection burden was far less sensitive to these parameters and was mainly sensitive to human recovery rate (*γ*_*h*_) (Fig. 7(a)-(b)).

## Discussion

We are interested in what drives arbovirus epidemics with Zika as a model and how to reduce the burden of these diseases, focusing on temperature and key parameters that correspond to existing or potential control methods (e.g., pesticides, reduced breeding habitats, vaccines, or treatment). We investigated temperature-dependent dynamic transmission models that incorporated recent empirical estimates of the relationships between temperature and Zika infection, transmission, and mosquito lifespan^1^. These dynamical models that can measure final epidemic size and account for temperature variation generate qualitatively different results than static *R*_0_ variables. Temperature had an overwhelmingly strong impact on both *R*_0_ and the final epidemic size (total infectious individuals, equivalent to area under the *I*_*h*_ epidemic curve), but the response was much more gradual and had a clear optimum for *R*_0_, while the final epidemic size responded as a threshold function (Fig. 2(i)-(j)). This is because, while epidemics have a higher peak at the max *R*_0_ (at optimal temperatures), the epidemics are longer at sub-optimal temperatures (lower *R*_0_). Thus, Zika virus is capable of spreading efficiently through the host population (high *I*_*h*_) across a broad range of temperatures for which *R*_0_ > 1, spanning from 17-37° C in constant environments (Fig. 6) (a)^9^. This is broadly consistent with the high seroprevalence of Zika found in a number of countries^47, 48^. This suitable temperature region expanded and shifted toward cooler mean temperatures under seasonally varying environments (Fig. 6 (a)).

These results have two key implications. First, large epidemics can occur under realistic, seasonally varying, temperature environments even in regions where the mean temperature alone would be expected to suppress transmission, for example in a location with a mean of 15° C and a seasonal amplitude of 7° C. Second, temperature determines both upper and lower thresholds for whether or not epidemics are possible^9^. However, within the predicted suitable temperature range defined by *R*_0_, the final epidemic size is largely limited by the density of susceptible hosts (Figs. 2, 6 (a))^9^. More broadly, the results highlight the important principle that metrics of transmission (e.g., *R*_0_) have a nonlinear relationship with the human final epidemic size (total *I*_*h*_) and contribute distinct implications for our understanding of the transmission process.

Whether or not temperature affects the potential for disease control via vector control, reduction in host biting rate, vaccination, or drug administration is an important applied question for designing public health campaigns. Temperature did not strongly affect the impact of most control-related parameters on the final epidemic size when the models did not include seasonal variation. When the models included seasonal variation, the effectiveness of most control parameters depended on temperature. In all models, human vaccination rate required to control epidemics varied strongly with mean temperature (Figs. 3-6). Achieving herd immunity and thereby suppressing transmission via vaccination is more difficult when temperatures are highly suitable (20^0^-35^0^ C under constant temperatures or 15^0^-32^0^ C under varying temperatures; Figs. 3-6). By contrast, the effects of the human recovery rate (*γ*_*h*_) and the vector mortality rate (*µ*_*v*_) on *R*_0_ were sensitive to temperature, but their effects on the final epidemic size were not sensitive to temperature.

Similar to previous work on dengue^9^, our results show that Zika can invade and cause large outbreaks during the summer in seasonally varying environments with lower average temperatures, such as temperate regions of the U.S., Europe, and Asia. This implies that differences in the size of epidemics in tropical versus temperate locations occur not just because of differences in temperature (and its impacts on *R*_0_) but also because of differences in vector breeding habitat availability, humidity, human mosquito exposure, and other socio-ecological factors. Much of the globe—including regions in temperate, subtropical, and tropical climates—is already suitable for Zika transmission for all or part of the year, and climate change is likely to expand this suitability geographically and seasonally (Ryan et al., in review). However, processes that increase the density of susceptible human populations and their exposure to mosquitoes, including urbanization and urban poverty, human population growth, and the growth and geographic expansion of vector populations, are likely to expand the burden of Zika even more dramatically in the future.

There are several limitations of such a modeling study. First, the parameters are determined by a combination of lab-based estimates as well as from literature on dengue, instead of being fitted to empirical data on the spatio-temporal dynamics of Zika from the field. Such divergent approaches can generate different parameter estimates. Further, the projections of the model on potential geographic distribution of Zika are based on average of temperatures by country and season with constant parameters. In reality, there is substantial heterogeneity of temperature and parameters over time and space, which have important implications for disease dynamics. For this reason, further investigation of Zika models that are calibrated from field based dynamics will be valuable for a fuller understanding of effects of temperature variation on Zika control.

## Conclusion

The unexpected emergence and global expansion of Zika in 2015-2017 and its association with Zika congenital syndrome and Guillain-Barre syndrome revealed once again how poorly prepared the global community is for the looming and expanding threat of vector-borne diseases. Given the recent history of *Aedes aegypti*-transmitted viruses, including dengue, chikungunya, and Zika, rapidly expanding worldwide and the challenges of controlling these epidemics without specific vaccines or drugs, understanding the ecological drivers of transmission and their effects on potential disease control tools is crucial for improving preparedness for future vector-borne disease emergence. If a Zika vaccine becomes available, then the precisely defined temperature thresholds for large epidemics predicted in our model imply that vaccination targets should be set based on climate. By contrast, because other potential interventions that would reduce vector population sizes, biting rates, and human recovery rates act more independently of temperature, targets could be set based on other socio-ecological factors in a given epidemic setting. This dynamic temperature-dependent modeling framework, which depends most strongly on vector and host parameters that are virus-independent, may be a useful first step for responding to future *Aedes*-borne disease epidemics.

## Acknowledgements

This project was supported by the National Science Foundation (Grant #1640780). CNN acknowledges the support of the Simons Foundation (Award #627346). EAM was supported by grants from the NIH (R35GM133439) and NSF (DEB-1518681).

## Author contributions statement

CNN, EAM, CCM, and MHB conceived of the study. TB, LRD, and CCM, conducted laboratory study. CNN conducted mathematical analysis. SJR conducted geographic analysis. CNN, EAM, CCM, SJR, and MHB wrote the manuscript. All authors reviewed the manuscript.

## Additional information

Competing interests: I declare that the authors have no competing interests or other interests that might be perceived to influence the results and/or discussion reported in this paper.

